# Panton-Valentine leucocidin is the key determinant of *Staphylococcus aureus* pyomyositis in a bacterial genome-wide association study

**DOI:** 10.1101/430538

**Authors:** Bernadette C Young, Sarah G Earle, Sona Soeng, Poda Sar, Varun Kumar, Songly Hor, Vuthy Sar, Rachel Bousfield, Nicholas D Sanderson, Leanne Barker, Nicole Stoesser, Katherine RW Emary, Christopher M Parry, Emma K Nickerson, Paul Turner, Rory Bowden, Derrick Crook, David Wyllie, Nicholas PJ Day, Daniel J Wilson, Catrin E Moore

## Abstract

Pyomyositis is a severe bacterial infection of skeletal muscle, commonly affecting children in tropical regions and predominantly caused by *Staphylococcus aureus*. To understand the contribution of bacterial genomic factors to pyomyositis, we conducted a genome-wide association study of *S. aureus* cultured from 101 children with pyomyositis and 417 children with asymptomatic nasal carriage attending the Angkor Hospital for Children in Cambodia. We found a strong relationship between bacterial genetic variation and pyomyositis, with estimated heritability 63.8% (95% CI 49.2-78.4%). The presence of the Panton-Valentine leucocidin (PVL) locus increased the odds of pyomyositis 130-fold (*p* =10^-17.9^). The signal of association mapped both to the PVL-coding sequence and the sequence immediately upstream. Together these regions explained > 99.9% of heritability. Our results establish staphylococcal pyomyositis, like tetanus and diphtheria, as critically dependent on expression of a single toxin and demonstrate the potential for association studies to identify specific bacterial genes promoting severe human disease.

## Introduction

Microbial genome sequencing and bacterial genome-wide association studies present new opportunities to discover bacterial genes involved in the pathogenesis of serious infections.^1–6^ Pyomyositis is a severe infection of skeletal muscle most commonly seen in children in the tropics.^7–9^ In up to 90% of cases it is caused by a single bacterial pathogen, *Staphylococcus aureus*.^7–10^ There is evidence that some *S. aureus* strains have heightened propensity to cause pyomyositis – the incidence in the USA doubled during an epidemic of community-associated methicillin resistant *S. aureus* (CA-MRSA)^11^ – but molecular genetic investigation of *S. aureus* from pyomyositis has been limited.^12^

Panton-Valentine leucocidin (PVL), a well-known staphylococcal toxin causing purulent skin infections and found in epidemics caused by CA-MRSA, has been implicated in pyomyositis, pneumonia and other *S. aureus* disease manifestations, but its role is strongly disputed.^13–16^ PVL is a bipartite pore-forming toxin comprising the co-expressed LukF-PV and LukS-PV proteins,^17,18^ is encoded by *lukSF-PV*, which is usually carried on bacteriophages^13,17^ which facilitates *lukSF-PV* exchange between lineages.^19^ Although small case series testing for candidate genes have reported a high prevalence of PVL among pyomyositis-causing *S. aureus*, ^11,20,21^ a detailed meta-analysis found no evidence for an increased rate of musculoskeletal infection or other invasive disease in PVL-positive bacteria *versus* controls.^13^ These conflicting results may reflect insufficiently powered studies. However, candidate gene studies may also miss important variation elsewhere in the genome: a study claiming a critical role for PVL in the causation of severe pneumonia^15^ was later found to have overlooked mutations in key regulatory genes, capable of producing the virulent behaviour that had been attributed to PVL by the original study.^16^ Thus, based on the current evidence, opinion is divided as to whether PVL is an important virulence factor in pyomyositis, or merely an epiphenomenon, carried by bacteria alongside unidentified genetic determinants.^22,23^

Genome-wide association studies (GWAS) offer a means to screen entire bacterial genomes to discover genes and genetic variants associated with disease risk. They are particularly appealing because they enable the investigation of traits not readily studied in the laboratory, and do not require the nomination of specific candidate genes.^5^ Proof-of-principle GWAS in bacteria have already demonstrated the ability to successfully rediscover known antimicrobial resistance (AMR) determinants.^2,3,4^ However, AMR is under extraordinarily intense selection in bacteria, and it remains to be seen whether GWAS can overcome the typically strong linkage disequilibrium in bacterial populations to precisely pinpoint genes and genetic variants underlying the propensity to cause human infection.

## Results

To understand the bacterial genetic basis of pyomyositis, we sampled and whole-genome sequenced *S. aureus* from 101 pyomyositis infections and 417 asymptomatic nasal carriage episodes in 518 children attending Angkor Hospital for Children in Siem Reap, Cambodia between 2008 and 2012 (Table S1). As expected of *S. aureus* epidemiology, we observed representatives of multiple globally common lineages in Cambodia, together with some globally less common lineages at high frequency, in particular clonal complex (CC) 121, identified by Multi-locus sequence type (MLST). Lineage composition appeared stable over time, with no major changes in lineage frequency (Fig. S1).

In our study, some *S. aureus* lineages were strongly overrepresented among cases of pyomyositis compared to their frequency among asymptomatic, nasally-carried controls over the same time period. Notably, 86/101 (85%) of pyomyositis cases were caused by CC-121 bacteria, whereas no pyomyositis cases were caused by the next two most commonly carried lineages, sequence type (ST)-834 and CC-45 (Fig. 1). We estimated the overall heritability of case/control status to be 63.8% (95% CI 49.2-78.4%) in the sample, reflecting the strong relationship between bacterial genetic variation and case/control status. We used *bugwas*^6^ to decompose this heritability into the principal components (PCs) of bacterial genetic variation. PC 1, which distinguished CC-121 (the most common pyomyositis lineage) from ST-834 (which was only found in carriage), showed the strongest association with case /control status (*p* = 10^-29.6^, Wald test). The next strongest were PC 20, which differentiated a sub-lineage of CC-121 within which no cases were seen (*p* = 10^-13.9^), and PC 2, which distinguished CC-45 from the rest of the species (*p* = 10^-4.9^).

**Figure 1.**
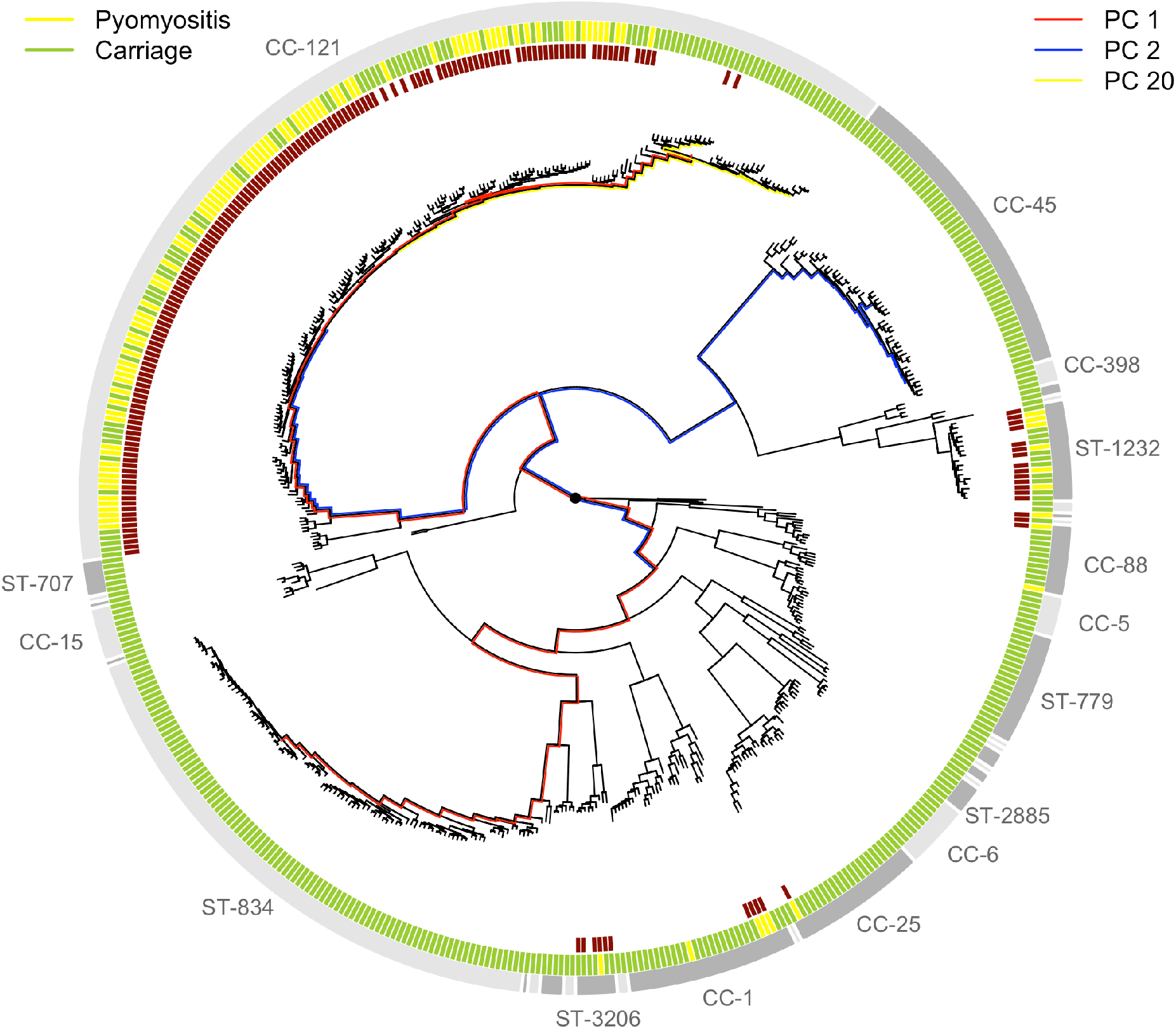
Phylogeny of *S. aureus* cultured from children in Cambodia shows strong strain-to-strain variation in pyomyositis prevalence. The phylogeny was estimated by maximum likelihood from SNPs mapping to the USA300 FPR3757 reference genome. Multi-locus sequence type (ST) or clonal complex (CC) groups are shown (outer gray ring). Case/control status is marked in the middle ring: pyomyositis (gold, n = 101) or nasal carriage (green, n = 417). Branches of the phylogeny that correspond to the three principal components (PCs) significantly associated with case/control status (PCs 1, 2 and 20) are marked in red, blue and yellow, respectively. Branch lengths are square root transformed to aid visualization. The presence of the kmers most strongly associated with pyomyositis is indicated by red blocks in the inner ring

We conducted a GWAS to identify bacterial genetic variants associated with pyomyositis, controlling for differences in pyomyositis prevalence between *S. aureus* lineages. We used a kmer-based approach^1^ in which every variably present 31bp DNA sequence observed among the 518 genomes was tested for association with pyomyositis *versus* asymptomatic nasal carriage, controlling for population structure using GEMMA.^24^ These kmers captured bacterial genetic variation including single nucleotide polymorphisms (SNPs), insertions or deletions (indels), and presence or absence of entire accessory genes. We found 10.7 million unique kmers variably present across the bacterial genomes. In total, 9,175 kmers were significantly associated with case/control status after correction for multiple testing (10^-6.8^ ≤ *p* ≤ 10^-21.4^; Fig 2A). The vast majority of these kmers (8,993/9,175; 98.0%) localised to a 45.7kb region spanning an integrated prophage with 95% nucleotide similarity to φSLT (Fig 2B). Most significant kmers, (9,173/9,175; 99.98%) were found more frequently in pyomyositis, with odds ratios (OR) ranging from 2.7 to 139.8, indicating they were associated with increased risk of disease. The bacteriophage φSLT was thus strongly associated with pyomyositis.

**Figure 2.**
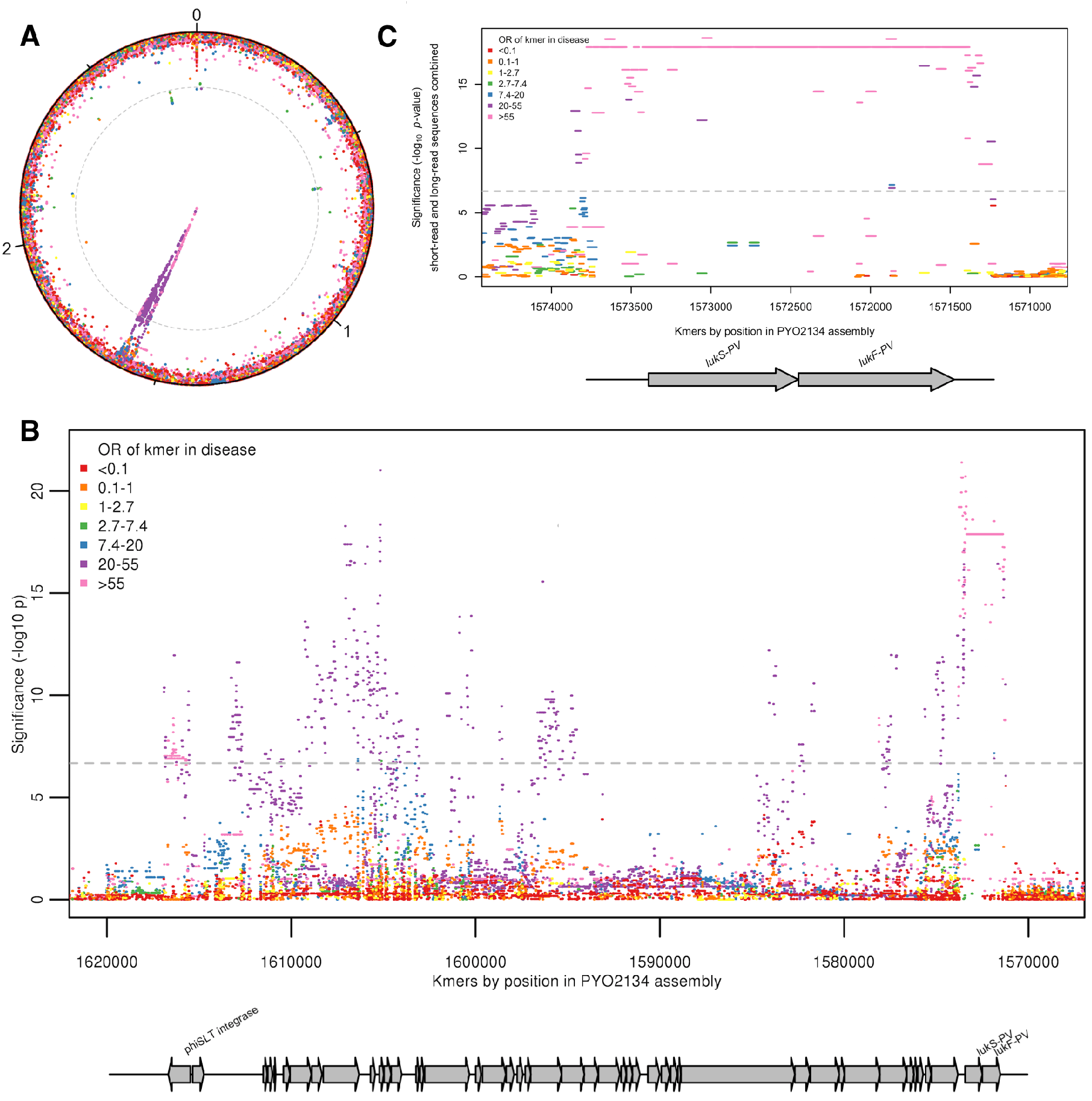
Kmers associated with pyomyositis. **(A)** All kmers (n = 10,744,013) were mapped to the genome assembly of one CC121 pyomyositis bacterium (PYO2134). Each point represents a kmer, plotted by the mapped location and the significance of the association with disease (-log_10_ *p* value). Kmers are coloured by the odds ratio (OR) of kmer presence for disease risk. A Bonferroni-adjusted threshold for significance is plotted in grey **(B)** The region between 1.57–1.62 MB in greater detail. Gray arrows depict coding sequences, determined by homology to USA300 FPR3757. **(C)** Associations for kmers mapping to region 1,571 – 1,574kB is plotted. Kmer presence determined from hybrid assembly using short and long-read data for assembly. Gray arrows depict coding sequences, determined by homology to USA300 FPR3757.

We were able to fine-map the signal of association within jSLT to the *lukS-PV* and *lukF-PV* cargo genes. These genes encode the subunits of PVL, which multimerise into a pore-forming toxin capable of rapidly lysing the membranes of human neutrophils, the first line of defense against *S. aureus*.^17,25^ 1630 kmers tagging the presence of the *lukSF-PV* coding sequences (CDS) were highly significantly associated with disease, being present in 98/101 (97%) pyomyositis cases and 84/417 (20%) carriage controls (OR 129.5, *p* = 10^-17.9^). Kmers tagging variation in the 389bp region immediately upstream of the CDS were also strongly associated with disease (*p* = 10^-21.4^). The most significant of these kmers were co-present with the CDS in the same cases (98/101, 97.0%), but present in fewer controls (79/417, 18.9%), producing an OR of 140.

Closer examination of this ∼400bp upstream region in genomes assembled from short-read Illumina sequencing showed that assembly of the region was problematic, with breaks or gaps in the assembly (Fig. S2). To improve the accuracy of this region of the assembled genomes we performed long-read Oxford Nanopore sequencing on the 37 genomes with incomplete or discontinuous assembly upstream of the PVL CDS. By integrating long-read and short-read data we were able to assemble a single contig spanning this region in all isolates (Fig. S3). When these improved assemblies were introduced, the signal of association upstream of the PVL CDS was no more significant than within the CDS (Fig 2C). Therefore, the presence of genomic sequence spanning the PVL toxin-coding sequences and the upstream, presumed regulatory, region exhibited the strongest association with pyomyositis in the *S. aureus* genome.

Presence or absence of the PVL region accounted for the differences in pyomyositis rates between lineages. It was common in pyomyositis-associated lineages including CC-121 and absent from non-pyomyositis-associated lineages including ST-834 and CC-45 (Fig. 1), explaining 99.9% of observed heritability in case-control status. It was infrequent in the non-pyomyositis-associated sub-lineage of CC-121 (2/36, 5.6%), and sporadically present in pyomyositis cases in otherwise non-pyomyositis-associated, PVL-negative strains CC-1 and CC-88. Its absence from only three cases (in CC-88, CC-1 and CC-121) suggested that the PVL region approached necessity for development of pyomyositis in Cambodian children, while its presence in 20% of controls indicated that PVL-associated pyomyositis is incompletely penetrant, i.e. presence of the PVL region does not always lead to disease.

## Discussion

In this study we found a strong association between pyomyositis, a highly distinctive tropical infection of skeletal muscle in children, and Panton-Valentine leukocidin, a bacterial toxin commonly carried by bacteriophages. We found that a single coding region together with the upstream sequence are all but necessary for the development of pyomyositis: its sporadic presence is associated with pyomyositis in otherwise low-frequency strains, and its absence is associated with asymptomatic carriage in a high propensity strain. The locally common PVL-positive CC-121 lineage contributes most strongly to the prevalence of pyomyositis in Cambodian children.

While PVL has long been thought an important *S. aureus* virulence factor,^25,26,27^ its role in invasive disease has been controversial,^22,23^ with conflicting results in case-control studies and an absence of supporting evidence on meta-analysis.^13^ In previous studies the PVL positive USA300 lineage was associated with musculoskeletal infection (both pyomyositis and osteomyelitis), however in these studies almost all these infections were caused by the USA300 strain, so the role of PVL was almost completely confounded by both methicillin resistance and strain background.^11,26^ In our study, this confounding is broken down by the movement of PVL on mobile genetic elements (MGEs). Here, despite the emergence of CA-MRSA in carriage in the same population,^28^ all the pyomyositis cases were MSSA. By applying GWAS methods to a well-powered cohort, our study resolves the controversy around PVL and pyomyositis, demonstrating strong heritability which localises to a single region, even when the full bacterial genome is considered. This study offers empirical evidence that, in addition to elucidating phenotypes under strong selection (such as in antimicrobial resistance)^2,3,4^, bacterial GWAS can pinpoint variants when MGEs act to unravel linkage disequilibrium.

These results offer a novel prospect for disease prevention: they establish staphylococcal pyomyositis as a disease whose pathogenesis depends critically on expression of a single toxin. This property is shared by toxin-driven, vaccine-preventable diseases such as tetanus and diphtheria. Therefore, vaccines that generate neutralising anti-toxin antibodies against PVL^29^ may protect human populations against this common tropical disease. They also raise the hypothesis that antibiotics which decrease toxin expression, and have been recommended in some PVL-associated infections,^30^ may offer specific clinical benefit in treating pyomyositis. More generally, our study provides an example of how microbial GWAS can be used to elucidate the pathogenesis of bacterial infections and identify specific virulence genes associated with human disease.

## Materials and Methods

### Patient sample collection

We retrospectively identified pyomyositis cases from the Angkor Hospital for Children in Siem Reap, Cambodia, between January 2007 and November 2011. We screened all attendances in children (≤16 years) using clinical coding (ICD-10 code M60 (myositis)) and isolation of *S. aureus* from skeletal muscle abscess pus. We reviewed clinical notes to confirm a clinical diagnosis of pyomyositis was made by the medical staff, and bacterial strains cultured by routine clinical microbiology laboratory were retrieved from the local microbiology biobank. 106 clinical episodes of pyomyositis were identified, in 101 individuals, and we included the earliest episode from each individual.

We identified *S. aureus* nasal colonisation from two cohort studies undertaken at Angkor Hospital for Children. The first were selected from a collection characterising nasal colonization in the region between September and October, 2008, which has previously been described using multi-locus sequence typing.^28^ The swabs had been saved at -80°C since the study, these samples were reexamined for the presence of *S. aureus* using selective agar, confirmed using Staphaurex (Remel, Lenexa, USA) and the DNAse agar test (Oxoid, Hampshire, UK). Antimicrobial susceptibility testing was performed according to the 2014 Clinical and Laboratory Standards Institute guidelines (M100-24).^31^

We undertook a second cohort study in 2012. Inclusion criteria were children (≤16 years) attending as an outpatient at Angkor Hospital for Children with informed consent. There were no exclusion criteria. Children were swabbed between the 2-7th July 2012, using sterile cotton tipped swabs pre-moistened (with phosphate buffered saline, PBS) using 3 full rotations of the swab within the anterior portion of each nostril with one swab being used for both nostrils, the ends were broken into bottles containing sterile PBS and kept cool until plated in the laboratory (within the hour). The swabs were plated onto Mannitol Salt agar to select for *S. aureus*. The M100–24 CLSI^31^ standards were followed for susceptibility testing and bacteria stored in tryptone soya broth and glycerol at -80°C.

We selected controls from carriers in these two cohorts using the excel randomization function: 222 of 519 from the 2008 cohort and 195 of 261 from the 2012 cohort.

### Ethical Framework

Approval for this study was provided by the AHC institutional review board and the Oxford Tropical Ethics Committee (507-12).

### Whole genome sequencing

For each bacterial culture, a single colony was sub-cultured and DNA was extracted from the sub-cultured plate using a mechanical lysis step (FastPrep; MPBiomedicals, Santa Ana, CA) followed by a commercial kit (QuickGene; Autogen Inc, Holliston, MA), and sequenced at the Wellcome Trust Centre for Human Genetics, Oxford on the Illumina (San Diego, California, USA) HiSeq 2500 platform, with paired-end reads 150 base pairs long.

A subset of samples were sequenced using long-read sequencing technology. We selected 37 isolates with incomplete assembly upstream of the PVL locus, 22 with ambiguous base calls in the assembly, and 15 where the region was assembled over 2 contigs. DNA was extracted using Genomic Tip 100/G (Qiagen, Manchester, UK) and DNA libraries prepared using Oxford Nanopore Technologies (ONT) SQK-LSK108 library kit (ONT, Oxford, UK) according to manufacturer instructions. These were then sequenced on ONT GridION device integrated with a FLO-MIN106 flow cell (ONT, Oxford, UK). ONT base calling was performed using Guppy v.1.6.

### Variant calling

For short-read sequencing we used Velvet^32^ v1.0.18 to assemble reads into contigs *de novo*. Velvet Optimiser v2.1.7 was used to choose the kmer lengths on a per sequence basis. The median kmer length was 123bp (IQR 119-123). To obtain multilocus sequence types we used BLAST^33^ to find the relevant loci, and looked up the nucleotide sequences in the online database at http://saureus.mlst.net/. Strains that shared 6 of 7 MLST loci were considered to be in the same Clonal Complex. Antibiotic sensitivity was predicted by interrogating the assemblies for a panel of resistance determinants as previously described.^34^

We used Stampy^35^ v1.0.22 to map reads against reference genomes (USA300_FPR3757, Genbank accession number CP000255.1).^3^6 Repetitive regions, defined by BLAST^3^3 comparison of the reference genome against itself, were masked prior to variant calling. Bases were called at each position using previously described quality filters.^37–39^

After filtering ONT reads with filtlong v.0.2.0 (with settings filtlong -- min_length 1000 -- keep_percent 90 --target_bases 500000000 --trim --split 500), hybrid assembly of short (Illumina) and long (ONT) reads were made, using Unicycler v0.4.5^40^ (default settings). The workflow for these assemblies is available at https://gitlab.com/ModernisingMedicalMicrobiology/MOHAWK)

### Reconstructing the phylogenetic tree

We constructed a maximum likelihood phylogeny of mapped genomes for visualization using RAxML^41^ assuming a general time reversible (GTR) model. To overcome a limitation in the presence of divergent sequences whereby RAxML fixes a minimum branch length that may be longer than a single substitution event, we fine-tuned the estimates of branch lengths using ClonalFrameML.^42^

### Kmer counting

We used a kmer-based approach to capture non-SNP variation.^1^ Using the *de novo* assembled genome, all unique 31 base haplotypes were counted using dsk^43^. If a kmer was found in the assembly it was counted present for that genome, otherwise it was treated as absent. This produced a set of 10,744,013 variably present kmers, with the presence or absence of each determined per isolate. We identified a median of 2,801,000 kmers per isolate, including variably present kmers and kmers common to all genomes (IQR 2,778,000–2,837,000).

### Calculating heritability

We used the Genome-wide Efficient Mixed Model Association tool (GEMMA^24^) to fit a univariate linear mixed model for association between a single phenotype (pyomyositis vs asymptomatic nasal carriage). We calculated the relatedness matrix from kmers, and used GEMMA to estimate the proportion of variance in phenotypes explained by genotypic diversity (i.e. heritability).

### Genome wide association testing of Kmers

We performed association testing using an R package bacterialGWAS (https://github.com/jessiewu/bacterialGWAS), which implements a published method for locus testing in bacterial GWAS.^3^ The association of each kmer on the phenotype was tested, controlling for the population structure and genetic background using a linear mixed model (LMM) implemented in GEMMA.^24^ The parameters of the linear mixed model were estimated by maximum likelihood and likelihood ratio test was performed against the null hypothesis (that each locus has no effect) using the software GEMMA.^24^ GEMMA was run using a minor allele frequency of 0 to include all SNPs. GEMMA was modified to output the ML log-likelihood under the null, and alternative and–log_10_ *p* values were calculated using R scripts in the bacterialGWAS package. Unadjusted odds ratios were reported because there was no residual heritability unexplained by the most significant kmers.

### Testing for lineage effects

We tested for associations between lineage and phenotype using an R package *bugwas* (available at https://github.com/sgearle/bugwas), which implements a published method for lineage testing in bacterial GWAS.^3^ We tested lineages using principal components. These were computed based on biallelic SNPs using the R function prcomp. To test the null hypothesis of no background effect of each principal component, we used a Wald test, which we compared against a *χ*^2^ distribution with one degree of freedom to obtain a *p* value.

### Kmer mapping

We used Bowtie^44^ to align all 31bp kmers from short-read sequencing were to a draft reference (the *de novo* assembly of a CC-121 pyomyosits strain PYO2134). Areas of homology between the draft reference and well-annotated reference strains were identified by aligning sequences with Mauve^45^. For all 31bp kmers with significant association with case-controls status, the likely origin of the kmer was determined by nucleotide sequence BLAST^33^ of the kmers against a database of all *S. aureus* sequences in Genbank.

### Joint short-read and long-read analysis

31bp kmers were counted for the 37 hybrid short-read and long-read assemblies using dsk^43^. The presence or absence of all Illumina (short-read) kmers that mapped to the two PVL toxin-coding sequences and the upstream intergenic region plus the surrounding 1kb were reassessed. For the 37 samples with hybrid assemblies, the presence/absence of these kmers was determined from the kmers counted from the hybrid assemblies. For all other samples, presence/absence was determined from the kmers counted from the short-read only assemblies. The new presence/absence patterns were tested for association with the phenotype controlling for population structure and genetic background using GEMMA^24^, using the same relatedness matrix as the original short-read analysis.

### Multiple testing correction

Multiple testing was accounted for by applying a Bonferroni correction;^46^ the individual locus effect of a variant (kmer or PC) was considered significant if its *P* value was smaller than *α*/*n*_p_, where we took *α* = 0.05 to be the genome-wide false-positive rate and *n*_p_ to be the number of kmers or PCs with unique phylogenetic patterns, that is, unique partitions of individuals according to allele membership. We identified 236627 unique kmer patterns and 518 PCs, giving thresholds of 2.1 x10-7and 9.7×10^-5^ respectively.

### Data availability

Sequence data has been submitted to Short Read Archive (Bioproject ID PRJNA418899).

## Acknowledgments

The authors would like to thank study participants. This study was funded by the Wellcome Trust (MORU Grants 089275/H/09/Z and 089275/Z/09/Z), and a University of Oxford Medical Research Fund awarded to C.E.M. (MRF/MT2015/2180). D.J.W. is a Sir Henry Dale Fellow, jointly funded by the Wellcome Trust and the Royal Society (Grant 101237/Z/13/Z). B.C.Y. is a Research Training Fellow funded by the Wellcome Trust (Grant 101611/Z/13/Z). D.H.W. was funded by the National Institute for Health Research (NIHR) Oxford Biomedical Research Centre (BRC) and the European Union’s Seventh Framework Programme under the grant agreement number 601783 (BELLEROPHON project). N.S. is funded by a Public Health England (PHE)/University of Oxford Clinical Lectureship. K.E. was funded by an academic clinical fellowship which was provided by the UK NIHR through the University of Oxford. This research was supported by Core funding to the Wellcome Centre for Human Genetics provided by the Wellcome Trust (090532/Z/09/Z). The views expressed are those of the author(s) and not necessarily those of the NHS, PHE, the NIHR or the Department of Health.

## Author contributions

NPJD, CMP and CEM designed the study. SS, PS, VK, SH, VS, RB, NS, KE, CMP, EN, PT & CEM collected bacterial samples and clinical data. CEM performed DNA extraction for Illumina sequencing. LB performed Nanopore sequencing. BCY, SGE and NDS performed bioinformatics on the study. BCY, SGE, DW, NPJD, DJW and CEM analysed the data. RB and DC assisted with interpretation of findings. BCY, DJW and CEM wrote the manuscript.

## Supplementary information

**Figure S1.**
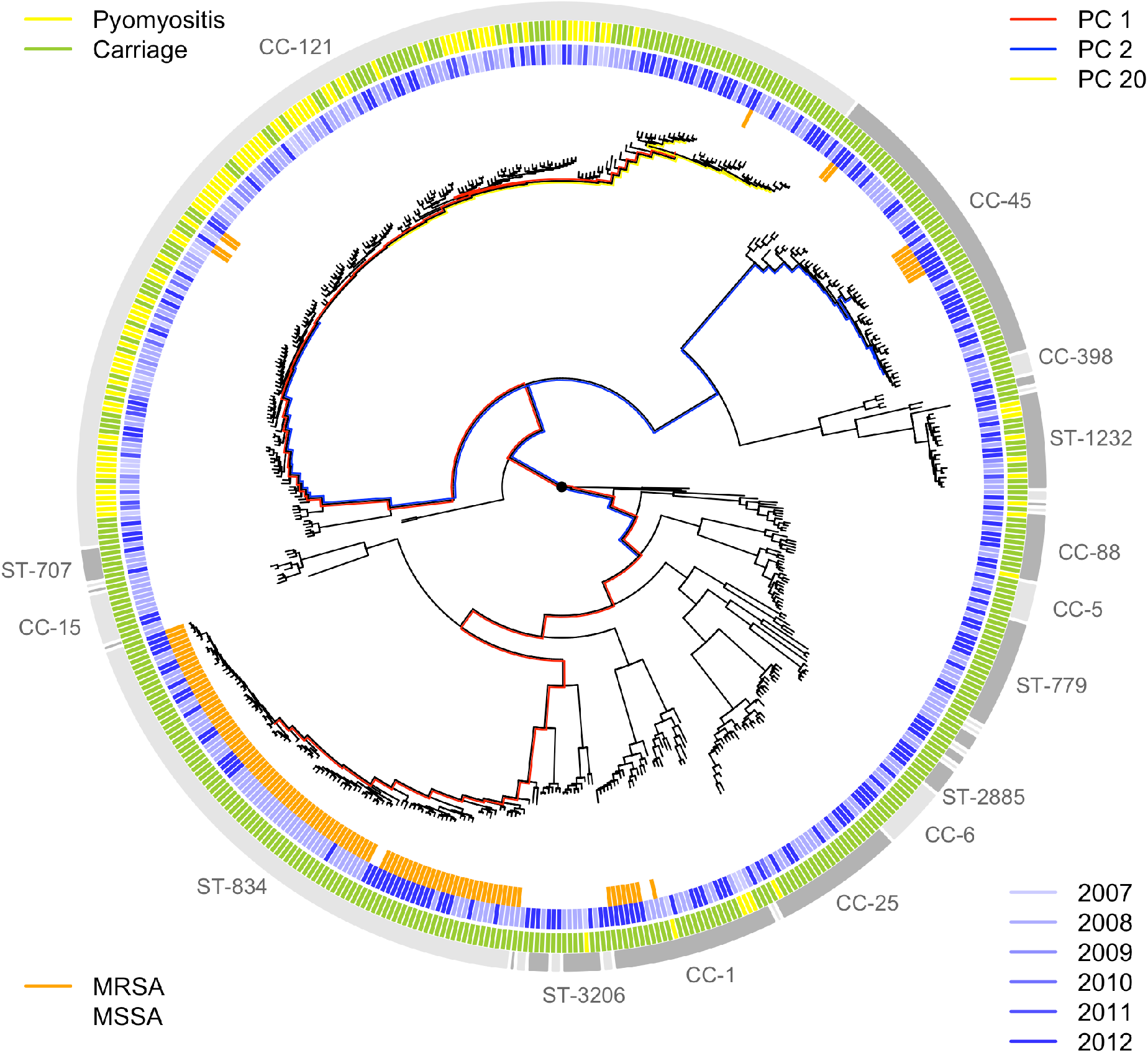
Sampling frequencies of the major strains were stable over time. The year of sampling (2007–2008, blue shaded lines) and MRSA status (orange lines) are illustrated around the phylogeny of the bacteria sampled from pyomyositis cases (gold lines) and asymptomatic carriage controls (green lines). The three PCs most significantly associated with case/control status are also shown (PCs 1, 2 and 20 by red, blue and yellow branches respectively).

**Figure S2.**
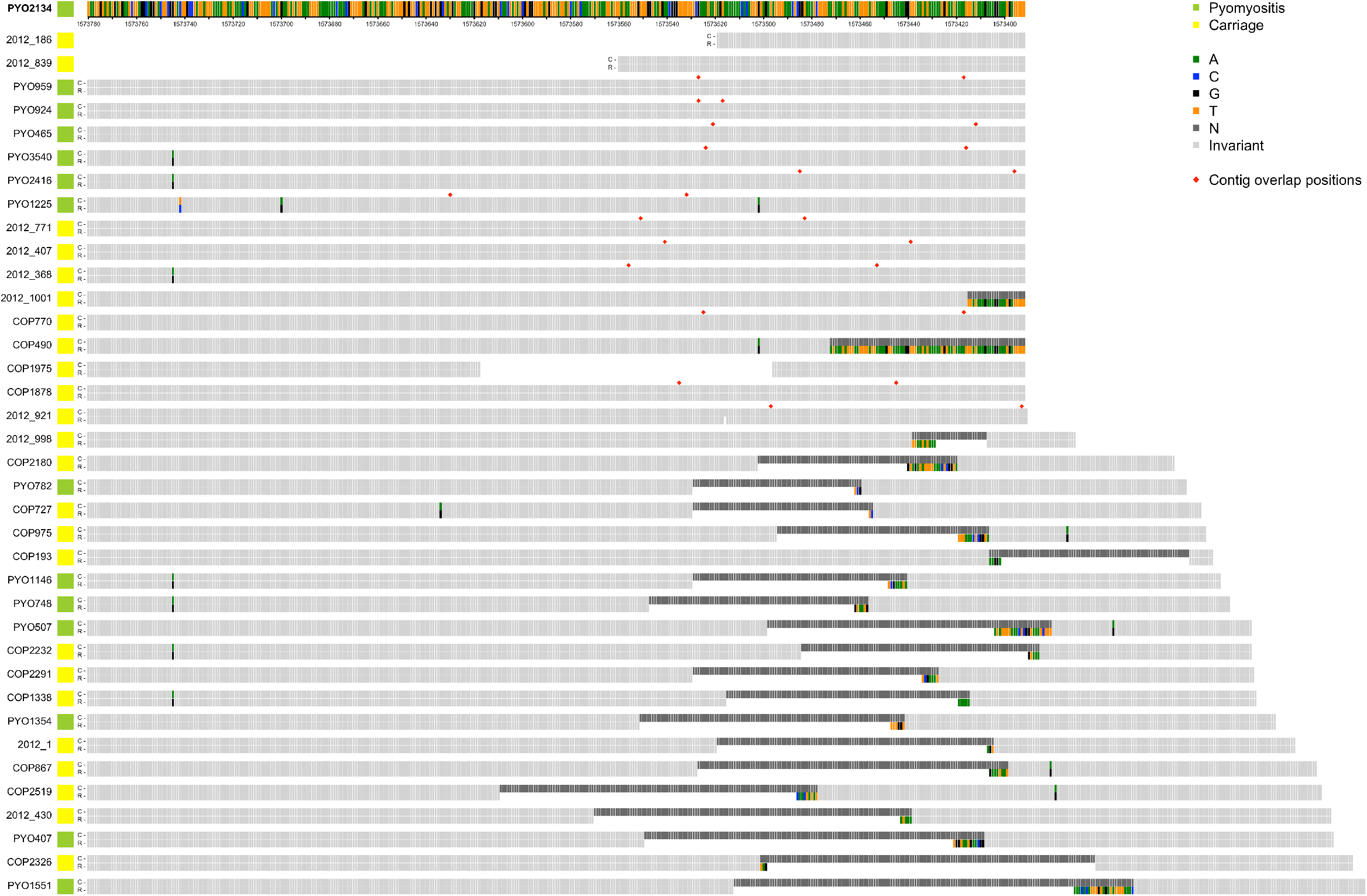
Alignments of reference genome PYO2134 assembly (R) with 37 *de novo* assemblies of Illumina short-read sequencing (C) which feature either ambiguities (Ns) or contig boundaries in the region 389bp upstream of PVL coding sequence. Contig boundaries, when overlapping, are marked with a red diamond. Ns in the assembly are marked in dark grey. Polymorphisms are colour-coded by base.

**Figure S3.**
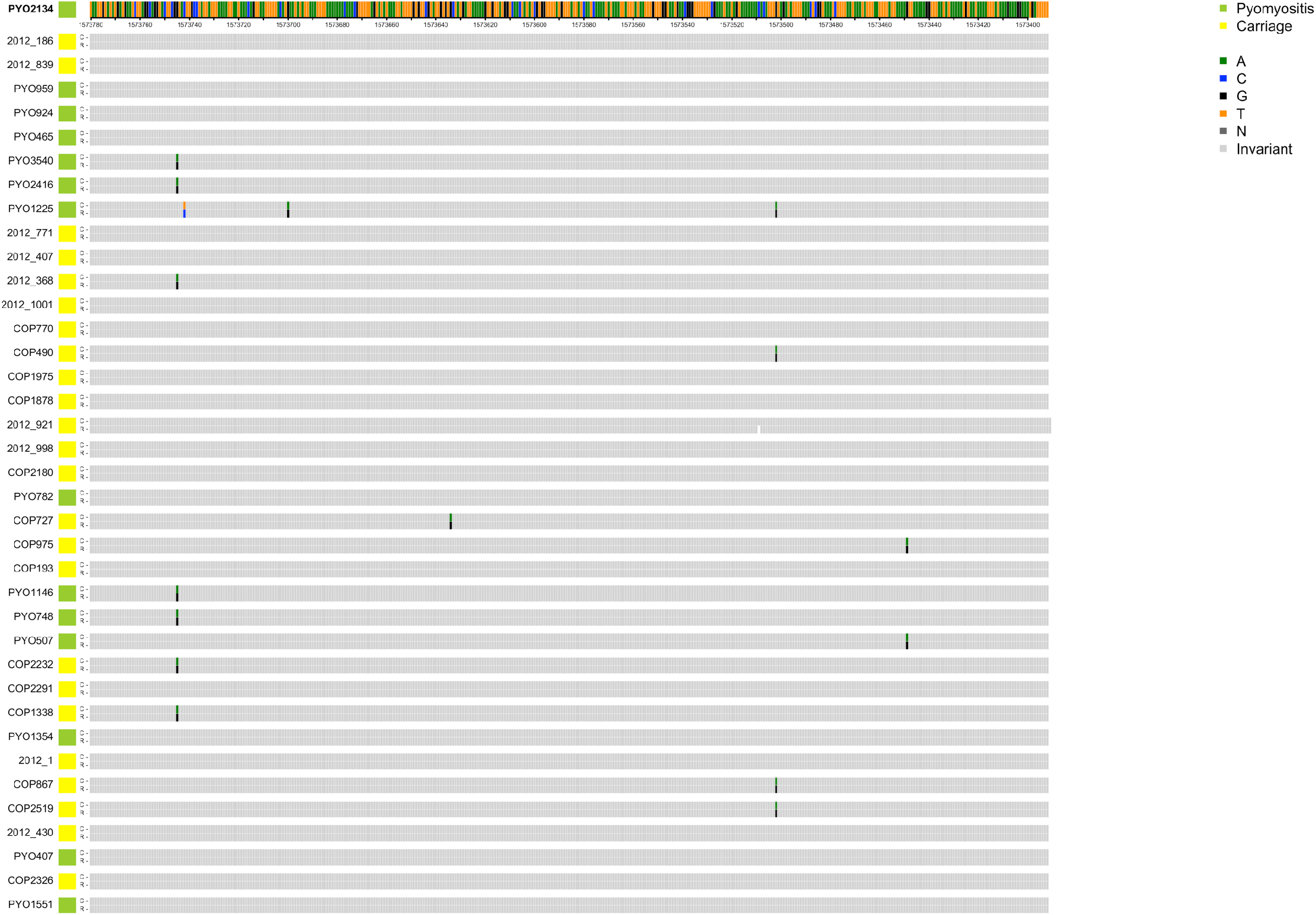
Alignments of reference genome PYO2134 assembly (R) with 37 *de novo* hybrid assemblies combining Oxford Nanopore long-read and Illumina short-read sequencing (C) which featured ambiguities (Ns) or hybrid assembly contig boundaries in the region 389bp upstream of PVL coding sequence in the Illumina short-read only assemblies. All ambiguities and contig boundaries are resolved in the hybrid assemblies.

**Table S1.** Isolates included in this study.

|  | Pyomyositis<br>(2007-2012) | Nasal carriage<br>(2008) | Nasal carriage<br>(2012) |
| --- | --- | --- | --- |
| Number of isolates | 101 | 222 | 195 |
| Age (med, IQR) | 7.8 years<br>(4.2-11.8) | 5.9 years<br>(2.5- 8.3) | 6.3 years<br>(4.1- 9.9) |
| Male (n (%)) | 66/97 (68%) | 122/221 (55.2%) | 105/195 (53.8%) |
| MRSA (n (%)) | 0 (0%) | 61 (27.5%) | 52 (26.7%) |

**Table S2:** All significant kmers from short read sequencing assembly, evidence of association, frequency and best match on blastn to all *S. aureus* sequences in Genbank.

